# Evaluation of machine learning models for proteoform retention and migration time prediction in top-down mass spectrometry

**DOI:** 10.1101/2021.10.31.466700

**Authors:** Wenrong Chen, Elijah N. McCool, Liangliang Sun, Yong Zang, Xia Ning, Xiaowen Liu

## Abstract

Reversed-phase liquid chromatography (RPLC) and capillary zone electrophoresis (CZE) are two popular proteoform separation methods in mass spectrometry (MS)-based top-down proteomics. The prediction of proteoform retention time in RPLC and migration time in CZE provides additional information that can increase the accuracy of proteoform identification and quantification. Whereas existing methods for retention and migration time prediction are mainly focused on peptides in bottom-up MS, there is still a lack of methods for the problem in top-down MS. We systematically evaluated 6 models for proteoform retention and/or migration time prediction in top-down MS and showed that the Prosit model achieved a high accuracy (*R*^2^ > 0.91) for proteoform retention time prediction and that the Prosit model and a fully connected neural network model obtained a high accuracy (*R*^2^ > 0.94) for proteoform migration time prediction.

## 1. Introduction

Top-down mass spectrometry (MS) is the method of choice for proteoform identification, characterization, and quantification ^1–3^. Many efforts have been made to increase proteoform identifications in proteome-wide studies based on top-down MS, and now thousands of proteoforms can be identified from a biological sample ^4^. Increasing proteome coverage is essential for many applications of top-down MS, such as disease biomarker identification ^5^. In top-down MS, the primary techniques for increasing proteome coverage are efficient proteoform separation methods and mass spectrometers with high speed, high resolution, and high accuracy.

Liquid chromatography (LC) and capillary zone electrophoresis (CZE) are two main techniques for protein separation in MS-based top-down proteomics ^6, 7^. In an LC experiment, proteins are separated based on their hydrophobicity, size, or other properties using various stationary phases in an LC column. There are many types of LC methods, such as reversed-phase liquid chromatography (RPLC) ^8^, size exclusion chromatography (SEC) ^9^, and ion exchange chromatography (IEC) ^10^. In top-down MS, RPLC is one of the most used methods due to its compatibility with popular, extensively developed, bottom-up proteomics platforms and high separation performance ^11, 12^.

In CZE-based separation, proteoforms are injected into a capillary filled with a background electrolyte on which an electric field is applied. Because proteoforms have different charges and hydrodynamic radii, they are separated by the migration time they need to move from one end to the other in the capillary, which is determined by their electrophoretic mobility ^13^. Many studies show that CZE is a highly efficient method for proteoform separation, with over a million theoretical plates achieved for some proteoforms ^14–16^.

Predicting proteoform retention time in RPLC-MS and migration time in CZE-MS can increase the accuracy of proteoform identification in top-down MS. When a mass spectrum is matched to an incorrect proteoform, there is often a large difference between the retention/migration time of the spectrum and the theoretical time of the proteoform. If the proteoform retention/migration time is accurately predicted, it can be used to filter out the proteoform identification whose theoretical retention/migration time does not match the empirical one, increasing proteoform identification accuracy.

Many methods have been proposed for retention time prediction in bottom-up MS^17^, which can be divided into three categories: library-based methods, index-based methods, and machine learning-based methods. In library-based methods, a library is built and maintained for the retention times of peptides identified from previous LC experiments, and retention time is predicted using the library. In index-based methods, retention coefficients of amino acids are first computed based on experimental data, and the retention time of a peptide is calculated as the sum of the retention coefficients of its amino acids. For example, SSRCalc ^18, 19^ produced high accuracy in retention time prediction using retention coefficients.

Machine learning-based methods achieved the best performance for retention time prediction in bottom-up MS. QSRR calculates and selects significant chemical descriptors of peptides and then performs a regression method to predict retention time ^20^. RTPredict ^21, 22^ and ELUDE ^23^ extract discriminant features of the amino acids in a peptide and predict retention time using support vectors machines (SVMs). GPTime ^24^ utilizes the features from ELUDE and a Gaussian process regression ^25^ model to obtain a high accuracy for retention time prediction. Recently, many deep learning models have been developed for retention time prediction in bottom-up MS,^26, 27^ which can be divided into three groups: convolutional neural network (CNN)-based models, such as DeepRT+ ^28^ and DeepLC ^29^, recurrent neural network (RNN)-based models, such as Prosit ^30^ and DeepMass ^31^, hybrid models with both CNN and RNN layers, such as DeepDIA ^32^ and AutoRT ^33^. These deep learning models significantly increased the accuracy of peptide retention time prediction to *R*^2^ > 0.95. For CZE migration time prediction, the size and charge of the peptide are two major features that affect the electrophoretic mobility and the migration time ^13, 34–37^. Semi-empirical models based on the two features produced an accuracy of *R*^2^ > 0.97 for electrophoretic mobility prediction on bottom-up MS data sets ^13^.

The retention/migration time prediction problem in bottom-up MS shares a high similarity with that in top-down MS, and the main difference is that proteoforms in top-down MS are longer than peptides in bottom-up MS. While many methods have been proposed for peptide retention/migration time prediction, only several studies have been done for proteoform retention/migration time prediction. The main reasons are that high-quality training data sets are lacking for the proteoform retention/migration prediction problem and that long proteoforms make the prediction problem more complicated.

Chen et al. extended the semi-empirical model for peptide migration time prediction to proteoform migration time prediction in top-down MS ^38^ and obtained an *R*^2^ = 0.98 on an *E. coli* CZE-MS data set. To the best of our knowledge, there have been no studies of the retention time prediction problem in top-down LC-MS.

We built one data set for proteoform retention time prediction and one data set for proteoform migration time prediction in top-down MS and evaluated the performance of 6 models including GPTime, fully connected neural network (FNN), Prosit, DeepRT+, DeepDIA and semi-empirical model for retention and/or migration time prediction on the data sets. Experimental results showed that the Prosit model achieved a high accuracy for retention time prediction (*R*^2^> 0.91) and that the Prosit model and FNN model obtained a high accuracy for migration time prediction (Prosit: *R*^2^ > 0.94; FNN: *R*^2^ > 0.94). We also assessed a transfer learning method in which peptides and their retention/migration times were employed for model pretraining and showed that transfer learning improved the prediction accuracy for some complex neural network models when the size of top-down MS training data was limited.

## 2. Methods

### 2.1 Top-down MS data sets

A top-down RPLC-MS/MS data set and a top-down CZE-MS/MS data set were used in this study. The RPLC-MS/MS data set was generated from ovarian tumor samples ^39^. A solid phase extraction column (360 μm o.d. × 150 μm i.d.) was used for trapping and desalting before separation. The separation process was performed with a dual-pump Waters nanoACQUITY UPLC system (Millford, Massachusetts) and a 50 cm length analytical column (360 μm o.d. × 100 μm i.d.) packed with 3 μm diameter C2 (Separation Methods Technology, Newark, Delaware). A 5 μL sample was loaded and separated with a 180-minute gradient from 99% solvent A to 35% solvent A with a 0.3 μL/min flow rate (A: 0.2% formic acid in water, B: 0.2% formic acid in acetonitrile). The separation system was coupled with a Velos Orbitrap Elite mass spectrometer (Thermo Fisher, San Jose, California). MS1 and MS/MS spectra were collected at a resolution of 240,000 and 120,000 at 200 m/z, respectively. The top 4 precursor ions in each MS1 spectrum were isolated with a 4 m/z window and fragmented with CID at a normalized collision energy of 35%. Ten technical replicates were generated for the same sample.

The CZE-MS/MS data set were obtained from SW480 colon cancer cells. Sample proteins were first separated by an SEC column into 6 fractions, and then each fraction was injected into an LPA (linear polyacrylamide) coated fused silica capillary (1m, 50 µm i.d., 360 µm o.d.) with 5% acetic acid as the background electrolyte. The electrospray voltage was 2-2.3kV and the separation voltage was 30 kV for 100 minutes. The CZE system was coupled with a Q-Exactive HF mass spectrometer (Thermo Fisher, San Jose, California). MS1 and HCD MS/MS spectra were collected at a resolution of 120,000 at 200 m/z. The top 5 precursor ions in each MS1 spectrum were analyzed using HCD MS/MS. Three technical replicates were obtained for each fraction, and only the first replicate was used in this study.

### 2.2 Proteoform identification

All raw MS files were converted to centroided mzML files using msconvert in ProteoWizard ^40^. TopFD (version 1.4.0) ^41^ was employed to deconvolute the centroided mass spectra to neutral monoisotopic masses of precursor and fragment ions. The deconvoluted MS/MS spectra were searched against the corresponding Uniprot proteome database (version Oct 23, 2019) for proteoform identification using TopPIC (version 1.4.0) ^41^. In database search, the error tolerance for precursor and fragment masses was set to 15 parts-per-million (ppm), and unknown mass shifts were not allowed. Cysteine carbamidomethylation was specified as a fixed modification for the SW480 data set, and no fixed modifications were set for the ovarian tumor data set. Proteoform-spectrum-matches (PrSMs) reported by database search were filtered with a stringent E-value cutoff of 10^−5^ to remove low confidence ones. These PrSMs were further clustered by merging PrSMs into the same cluster if the proteoforms of the PrSMs were from the same protein and the difference of their precursor masses was < 1.2 Da. The PrSM with the best E-value in each cluster was reported, and PrSMs with N-terminal acetylation were filtered out. Details of the parameter settings of TopPIC are given in Table S1 in the supplemental material. TopFD reported a retention or migration time for each identified proteoform, which was the apex time of the RPLC or CZE profile of the proteoform in the LC-MS or CZE-MS map. The apex times were used as empirical retention/migration times of identified proteoforms, which were further normalized by dividing them by the separation time of the experiment.

### 2.3 Machine learning models

Five machine learning models were assessed for predicting retention time in top-down RPLC-MS: the model in GPTime ^24^, an FNN model, DeepRT+ ^28^, Prosit ^30^, and DeepDIA ^32^. The last four models and a semi-empirical function ^38^ were also evaluated for predicting migration time in top-down CZE-MS. All the models were implemented in Python. The FNN and DeepRT+ models were implemented using the Pytorch package ^42^, and the Prosit and DeepDIA models using the Keras package ^43^ with the TensorFlow backend.

#### 2.3.1 GPTime model for retention time prediction

The model in GPTime with 62 features ^23, 24^ was used for proteoform retention time prediction in top-down MS. The first feature was the proteoform length and the second was the volume computed as the sum of the bulkiness indexes ^44^ of all amino acid residues in the proteoform. The other 60 features were computed for the 20 standard amino acids. Each of the 20 amino acids was represented by three features: the hydrophobicity index ^45^, the number of occurrences, and a retention index computed based on a linear regression model using training data ^23^. Gaussian process regression with the radial basis function (RBF) kernel was used for proteoform retention time prediction ^25^.

#### 2.3.2 A semi-empirical model for migration time prediction

The semi-empirical model in ref. ^38^ predicted proteoform migration time in CZE-MS using the molecular mass *M* and charge *Z* of the proteoform. The molecular mass was included to predict the size of the proteoform. The charge was estimated as the total number of positively charged amino acid residues (R, H, K, and the N-terminus) in the proteoform ^13^. The electrophoretic mobility of the proteoform was predicted as 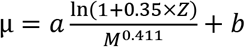, where *a* and *b* are two parameters related to the CZE settings ^38^. The electrophoretic mobility was then converted to its corresponding migration time using

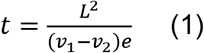

where *L* is the capillary length, *v*_1_ is the CZE separation voltage, and *v*_2_ is the electrospray voltage.

#### 2.3.3 Neural network models

An FNN model was built to predict retention or migration time in top-down MS, which contained an input layer, *k*(*k* = 1,2, or 3) fully connected hidden layers with dropout for regularization, and a fully connected output layer. The 62 features in the GPTime model were the input for retention time prediction. Five features were used for migration time prediction: the two features in the semi-empirical model and the numbers of D, E, N residues (see Results). For migration time prediction, we normalized proteoform masses by dividing them by 20,000 and normalized proteoform charges by dividing them by 20. The rectified linear unit (ReLU) activation function was used for the hidden layers, and the sigmoid function for the output layer. The model weights were initialized with a uniform distribution with zero mean and unit variance. The batch size was 8, the maximum training epochs was 12,000, the loss function was mean squared error (MSE), and the optimizer was the Adam algorithm with a learning rate of 10^−6^. The early stopping strategy was applied during the training process with a patience of 100. Various drop rates (0, 0.1, and 0.2) and node numbers (64, 128, 256, 512, 1024) for the hidden layers were tested (Table S2 in the supplementary material).

Three published neural network models were also assessed for predicting retention and migration time in top-down MS: CNN-based DeepRT+ ^28^, RNN-based Prosit ^30^, and a hybrid neural network model DeepDIA ^32^. In the three models, the loss function was MSE and the optimizer was Adam ^46^. The input of DeepRT+ and DeepDIA was the one-hot encoding of the amino acid sequence, and the input of Prosit was a sequence of 20 integers representing the amino acid sequence. Zero padding was added to the right end of the sequence to obtain the same length of 200, which was longer than the maximum proteoform length in the data sets. The learning rates for DeepRT+, Prosit, and DeepDIA were the default value 0.001.

In DeepRT+, the first two layers were convolutional ones, which were followed by two capsule layers connected by “dynamic routing” (Fig. S1 in the supplementary material). The root sum square of the output vector of the last capsule layer was reported as the predicted retention or migration time. Various hyperparameter settings were evaluated for the filter size and kernel size of the convolutional layers, the batch size, and the number of epochs (Table S3 in the supplementary material).

The Prosit model contained an embedding layer, a bidirectional GRU layer, a one-directional GRU layer, an attention layer, and two dense layers (Fig. S2 in the supplementary material). Hyperparameter settings, such as the unit number (64, 128, 256, and 512) in the GRU layers and the node number (64, 128, 256, and 512) in the dense layers, were tested to achieve the best prediction accuracy of Prosit (Table S4 in the supplementary material).

The DeepDIA model was composed of a convolutional layer, a max pooling layer, a bidirectional LSTM layer, and three dense layers (Fig. S3 in the supplementary material). A dropout layer with a rate of 0.5 was added between the LSTM and the first dense layer. We tuned the following hyperparameters of DeepDIA: the filter size and kernel size of the convolution layers, the number of units of the LSTM layer, and the number of features in the dense layers (Table S5 in the supplementary material).

### 2.4 Removing batch effects in migration time

Proteoform migration time in CZE-MS runs was affected by batch effect variations in these runs. The batch effects were removed with three steps. (1) Migration times were converted to their corresponding electrophoretic mobility values. (2) Batch effects in electrophoretic mobility were removed using the semi-empirical model and a method based on linear regression. (3) The electrophoretic mobility values with batch effect correction were converted back to migration times. Formula (1) in Section 2.3.2 was used for the conversion in the first and third steps. In the second step, electrophoretic mobility was predicted for each proteoform in a CZE-MS run using the semi-empirical model. Then a linear regression model *y* = *ax* + *b* was used to fit the experimental mobility *x* to the mobility *y* reported by the semi-empirical model in each fraction, where *a* and *b* are model parameters. For two CZE-MS runs, the electrophoretic mobility of proteoforms in the second run was mapped to that in the first run using the following method. Let *a*_1_ and *b*_1_ be the regression parameters for the first run, and *a*_2_ and *b*_2_ be the regression parameters for the second run. For a proteoform with mobility *x* in the second run, its mobility 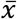 with batch effect correction satisfies the equation 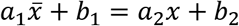, so the mobility with batch effect correction was computed as (*a*_2_*x* + *b*_2_ − *b*_1_)/*a*_1_.

### 2.5 Evaluation criteria

Three metrics were selected to evaluate the performance of the machine learning models: the MSE, *R*^2^, and Δ*t*_r95%_, where *R*^2^ measures the correlation between predicted and experimental time and Δ*t*_95%_, gives the minimal time window that explains 95% of the deviation between predicted and experimental time. The percentage of the Δ*t*_95%_ value compared with the overall elution/migration time was calculated, represented by Δ*t*_r95%_.

## 3. Results

### 3.1 Training and test data sets

TopPIC identified 610 proteoforms of 188 proteins from the first replicate of the RPLC-MS ovarian tumor (LC-OT) data. The LC-OT proteoforms were divided into 188 protein groups, which were then randomly split into a training set (131 protein groups with 437 proteoforms) and a test set (57 protein groups with 173 proteoforms) with a proteoform ratio of 7:3 approximately. Similarly, TopPIC reported from the first replicate of the CZE-MS/MS SW480 (CZE-SW480) data set 1230 proteoforms of 470 proteins, which were further randomly split by protein group into a training set of 878 proteoforms and a test set of 352 proteoforms.

### 3.2 Batch effect correction

The SW480 data set contained proteoforms identified from 6 SEC fractions of the sample, and the measured proteoform migration time was affected by variations in the CZE-MS runs (Fig. 1a). Because the fractions contain different proteoforms, time alignment ^47^ based on proteoform identifications is not a good method for batch effect correction. The semi-empirical model performed well in migration time prediction for single runs, but the variations in runs affected the prediction accuracy for the combined data (Fig. 1a). After batch effect correction (Methods), the *R*^2^ between experimental and predicted migration time were improved from 0.613 to 0.915 (Fig. 1b), showing that batch effect correction is an indispensable step for achieving high accuracy in proteoform migration time prediction.

**Figure 1.**
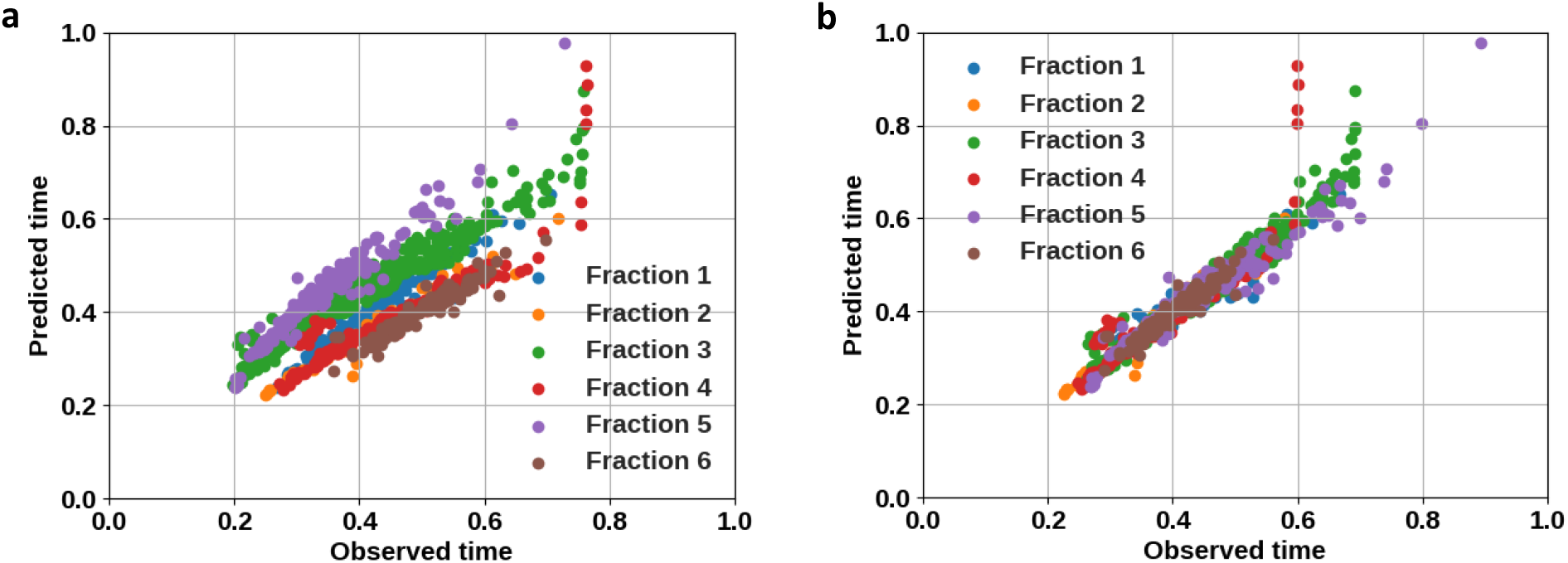
Batch error correction of migration time for the CZE-SW480 data with prefractionation. (a) Migration times predicted by the semi-empirical model are plotted against experimental migration times in 6 CZE-MS runs. The *R*^2^ between predicted and experimental migration time is 0.886 on average for single runs and 0.613 for the combined data of 6 runs. (b) The *R*^2^ between predicted and experimental migration time is improved to 0.915 for the combined data after batch error correction.

### 3.3 Retention time prediction

Hyperparameters were tuned for the FNN, DeepRT+, Prosit, and DeepDIA models using the LC-OT training set (437 proteoforms of 131 protein groups) with 5-fold cross validation. The 131 protein groups were divided into 5 folds so that each fold contained approximately the same number of proteoforms. The best hyperparameter settings for the 4 models are given in Tables S2-S5 in the supplementary material. Table 1 summarizes the prediction accuracy of the GPTime model and the four neural network models with the best hyperparameter settings on the LC-OT training set with 5-fold cross-validation. Prosit produced the best prediction accuracy (*R*^2^ = 0.906), and the conventional GPTime model outperformed FNN, DeepDIA, and DeepRT+. The low accuracy of FNN, DeepRT, and DeepDIA is possibly due to the small training data set. The performance of the 5 models was further compared by using the 7:3 training-test split of the LC-OT data set (Table S6 in the supplementary material), and the results were consistent with those on the LC-OT training set with 5-fold cross-validation.

**Table 1.** Benchmarking of 5 machine learning models for proteoform retention time prediction on the LC-OT training set with 5-fold cross validation.

| Model | $R^2$ | $\Delta t_{r95\%}$ | MSE |
| --- | --- | --- | --- |
| GPTIME | 0.885 | 0.311 | 0.00405 |
| FNN | 0.866 | 0.363 | 0.00479 |
| DeepRT+ | 0.731 | 0.531 | 0.00953 |
| Prosit | **0.906** | **0.293** | **0.00334** |
| DeepDIA | 0.789 | 0.424 | 0.00746 |

### 3.4 Migration time prediction

A total of 7 proteoform features in three groups were tested for proteoform migration time prediction: the molecular mass and the charge state (group 1), the numbers of D, E, and N residues (group 2), and the numbers of L and I residues (group 3). The high accuracy of the semi-empirical model ^38^ shows that the two features in group 1 are important for migration time prediction. D, E, and N residues (features in group 2) slightly influence the proteoform charge, and L and I residues (group 3 features) have the highest hydrophobicity indexes in CZE experiments.^48^ Four feature sets were compared with the FNN model with 2 hidden layers (256 nodes in each layer) on the CZE-SW480 training set with 5-fold cross validation: (1) group 1 only, (2) group 1 and group 2, (3) group 1 and group 3, and (4) all the features. The FNN model with the features in groups 1 and 2 obtained the best prediction accuracy *R*^2^= 0.959 (Table S7 in the supplementary material), showing that the features in group 2 provided additional information for migration time prediction.

Hyperparameter settings were tuned for the FNN, DeepRT+, Prosit, and DeepDIA models using the CZE-SW480 training set with 5-fold cross validation. The best hyperparameter settings of the models are given in Tables S2-S5 in the supplementary material. The hyperparameter settings selected for the models were not the same for RPLC and CZE, which is reasonable because the two separation methods are different. We tested the prediction accuracy of the semi-empirical model and 4 neural network models on two settings: the CZE-SW480 training set with 5-fold crossing validation and the 7:3 training-test split of the CZE-SW480 data set. Experimental results showed consistently that the performance of the Prosit and FNN models was comparable to the semi-empirical model and that the three models obtained better prediction accuracy than DeepRT+ and DeepDIA (Table 2 and Table S8 in the supplementary material). Whereas Prosit yielded a high prediction accuracy with a small training data set, DeepRT+ and DeepDIA suffered from the lack of large training data. The semi-empirical and FNN models reported high prediction accuracy with several proteoform features, indicating that it is possible to accurately predict proteoform migration time with simple models.

**Table 2.** Benchmarking of 5 machine learning models for proteoform migration time prediction on the CZE-SW480 training set with 5-fold cross validation.

| Model | $R^2$ | $\Delta t_{r95\%}$ | MSE |
| --- | --- | --- | --- |
| Semi-empirical | 0.934 | 0.194 | 0.00058 |
| FNN | **0.959** | **0.169** | **0.00034** |
| DeepRT+ | 0.726 | 0.415 | 0.00237 |
| Prosit | 0.943 | 0.186 | 0.00046 |
| DeepDIA | 0.874 | 0.277 | 0.00107 |

### 3.5 Transfer learning

Transfer learning ^49^ was adopted to address the problem that the training set was small in proteoform retention and migration time prediction. The DeepRT+ and Prosit models were first trained with a large data set of peptides and their retention/migration times identified by bottom-up MS, and then the weights in the models obtained from bottom-up MS data were used as initial weights in the training with top-down MS data. The hyperparameters of the models were the same as those in Tables S2-S5. The models for retention time prediction were pretrained using a bottom-up RPLC-MS/MS data set of 24 human cell lines and tissues including the HeLa cell line, muscle, and lung samples ^50^. X!Tandem ^51^ identified 146,587 peptides from the data set using database search, and the iRT Toolkit ^50^ reported normalized retention times of identified peptides. Detailed methods for peptide identification and retention time computation can be found in ref. ^50^. The DeepRT+ and Prosit models with and without pretraining were compared on the LC-OT data set with the 7:3 training-test split (Table 3). The transfer learning method significantly increased the accuracy of DeepRT+ from *R*^2^ 0.771 to 0.840 and the performance of Prosit from 0.860 to 0.914 (Fig. 2).

**Table 3.** DeepRT+ and Prosit with and without transfer learning are assessed on the LC-OT data set with a 7:3 training-test split for proteoform retention time prediction. The prediction accuracy on the test set is compared.

| Model | Without transfer learning |  |  | With transfer learning |  |  |
| --- | --- | --- | --- | --- | --- | --- |
| | $R^2$ | $\Delta t_{r95\%}$ | MSE | $R^2$ | $\Delta t_{r95\%}$ | MSE |
| DeepRT+ | 0.771 | 0.522 | 0.00857 | **0.840** | **0.378** | **0.00598** |
| Prosit | 0.860 | 0.486 | 0.00525 | **0.914** | **0.261** | **0.00321** |

**Figure 2.**
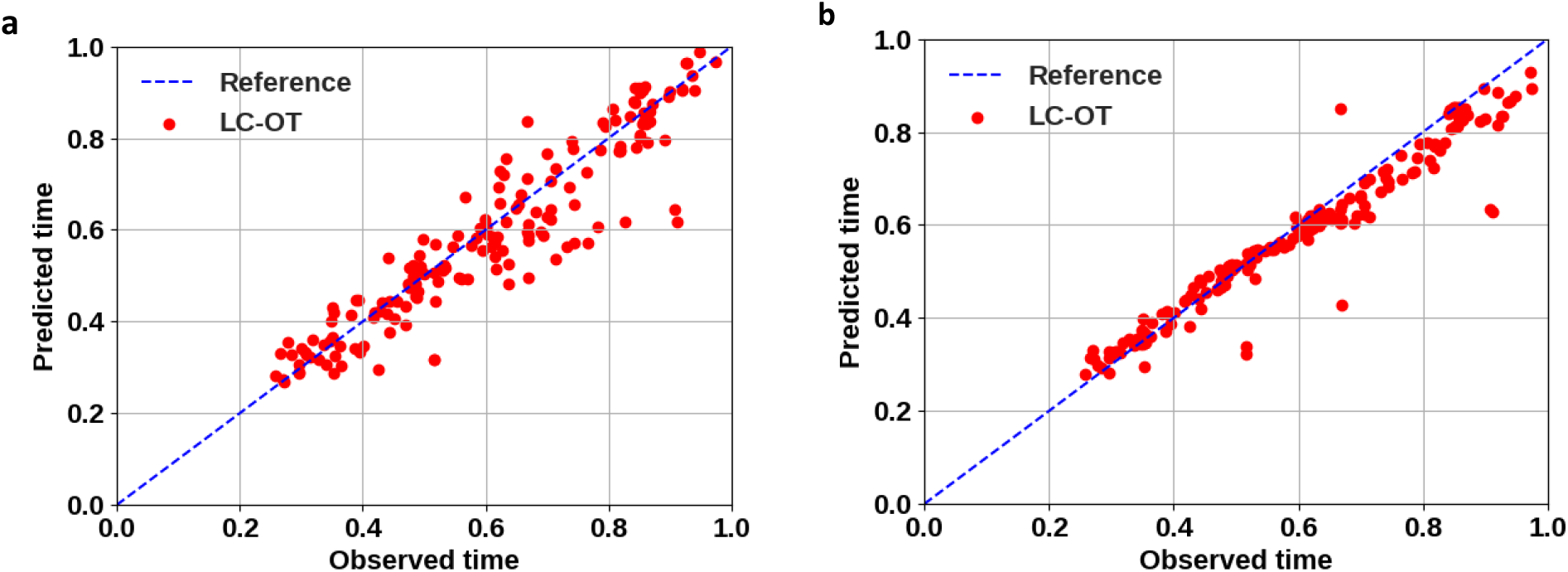
Comparison of the Prosit model with and without transfer learning on the LC-OT data. (a) The *R*^2^ of the Prosit model is 0.860 when it is trained with the LC-OT training set and tested on the LC-OT test set. (b) The *R*^2^ of the Prosit model is 0.914 when it is pretrained using a bottom-up data set of 146,587 peptides, trained with the LC-OT training set, and tested on the LC-OT test set.

The DeepRT+ and Prosit models for migration time prediction were pretrained using a bottom-up CZE-MS/MS data set of HeLa cells ^52^. The data set was generated from tryptic digests of proteins of HeLa cells and the spectra in the data set were analyzed by Mascot (version 2.2.4) in Proteome Discoverer 1.4 for peptide identification. We filtered out all identified peptides with PTMs or with a q-value > 0.001, resulting in 4,234 peptide identifications and their migration times. The two models with and without pretraining were compared on the CZE-SW480 data set with the 7:3 training-test split (Table 4). Pretraining with peptides improved the prediction accuracy for DeepRT+, but not for Prosit. The reason might be that pretraining did not provide additional useful information for the Prosit model, which can obtain a high accuracy for proteoform migration time prediction with a small training data set.

**Table 4.** DeepRT+ and Prosit with and without transfer learning are assessed on the CZE-SW480 data set with a 7:3 training-test split for proteoform migration time prediction. The prediction accuracy on the test set is compared.

| Models | Without transfer learning |  |  | With transfer learning |  |  |
| --- | --- | --- | --- | --- | --- | --- |
| | $R^2$ | $\Delta\text{tr}_{95\%}$ | MSE | $R^2$ | $\Delta\text{tr}_{95\%}$ | MSE |
| DeepRT+ | 0.750 | 0.291 | 0.00243 | **0.888** | **0.186** | **0.00108** |
| Prosit | **0.946** | **0.112** | **0.00052** | 0.930 | 0.141 | 0.00068 |

## 4. Discussion

The Prosit model designed for retention time prediction in bottom-up MS achieved high accuracy (*R*^2^ > 0.9) for the problem in top-down MS, demonstrating that it is not significantly affected by the long length of proteoforms and the limited training data set. The GRU and attention layers in Prosit are designed for processing long sentences, so it might be inheritably suitable for the proteoform retention/migration time prediction problem. The prediction accuracy of the DeepRT+ and DeepDIA models dropped significantly for the prediction problem in top-down MS compared with bottom-up MS. The reasons might be that the training data sets were too small to complex deep learning models and that the models are not suitable for processing long sequences.

The four neural network models reported similar prediction accuracy for retention and migration time prediction, indicating that these models may be used for many prediction problems in proteomics. With only several features, the semi-empirical and FNN models obtained high accuracy for migration time prediction, and most of the models reported a higher accuracy for migration time prediction than retention time prediction, showing that retention time prediction is more complicated than migration time prediction.

Because of the similarity between peptides and proteoforms, transfer learning, in which a model is pretrained on a large data set obtained from bottom-up MS, can improve prediction accuracy for proteoform migration and retention time prediction. But it did not increase the accuracy of the Prosit model for migration time prediction. The performance of transfer learning may depend on the model architecture and whether there is information that is transferable from the training data.

The study of the CZE-SW480 data with prefractionation reveals that the variations in CZE runs significantly affect experimental migration time and that batch effect correction is an indispensable step for accurate time prediction. Most of the variations in CZE runs can be removed by a regression-based method. The existence of the batch effect also complicates the applications of migration/retention time prediction models: A model trained on one data set needs to be adjusted or retrained before it is used on another data set.

Migration/retention time prediction has the potential to increase proteoform identifications in top-down MS. However, when the accuracy is not high enough, the improvement for proteoform identification is limited. When proteoforms lack MS/MS spectra or confident spectral identification, migration/retention time prediction may become more important for proteoform identification.

## 5. Conclusions

In this paper, we assessed several machine learning models for proteoform migration and retention time prediction in top-down MS. The Prosit model achieved high accuracy for proteoform migration and retention time prediction, and the FNN model outperformed other models in proteoform migration time prediction. Experimental results on transfer learning also showed its potential to increase prediction accuracy by using peptides identified from bottom-up MS. Yet the prediction accuracy achieved by the models is still not high enough to significantly increase proteoform identifications. Many important problems need to be further studied in this area. PTMs in proteoforms and batch effects compound the proteoform migration/retention time prediction problem, which will be our next research direction. Experiment settings in RPLC-MS and CZE-MS cause shifts in retention or migration time. It is a challenging problem to predict retention/migration time for experiments with different settings. In addition, a large data set is needed to further test and improve machine learning models for the proteoform retention/migration prediction problem.

## Acknowledgments

The research was funded by NIH through the grants R01GM118470 (Liu, Sun, and Ning), R01GM125991 (Sun and Liu), and R01CA247863 (Sun, Hummon, and Liu).

## Availability

The code is available at https://github.com/wenronchen/rt_prediction

## Notes

### Competing Interest Statement

The authors have declared no competing interest.

