## Supplemental Material for "Evaluation of machine learning models for proteoform retention and migration time prediction in top-down mass spectrometry"

[1]Department of BioHealth Informatics, Indiana University-Purdue University Indianapolis, Indianapolis, IN 46202, USA, [2]Department of Chemistry, Michigan State University, East Lansing, MI 48824, USA, [3]Department of Biostatics and Health Data Sciences, Indiana University School of Medicine, Indianapolis, IN, 46202, USA, [4]Department of Biomedical Informatics, The Ohio State University, Columbus, Ohio 43210, USA, [5]Department of Computer Science and Engineering, The Ohio State University, Columbus, Ohio 43210, USA, [6]Translational Data Analytics Institute, The Ohio State University, Columbus, Ohio 43210, USA, [7]Tulane Center for Biomedical Informatics and Genomics, Tulane University, New Orleans, LA, 70112, USA, [8]Deming Department of Medicine, Tulane University, New Orleans, LA 70112, USA

**Table S1.** Parameter settings of TopPIC

| Parameter | Value |
|---|---|
| Number of combined spectra | 1 |
| Fragmentation method | FILE |
| Search type | TARGET+DECOY |
| Fixed modifications | None/C57 |
| Use TopFD feature file | TRUE |
| Maximum number of unexpected modifications | 0 |
| Error tolerance | 15 ppm |
| Spectrum-level cutoff type | FDR |
| Spectrum-level cutoff value | 0.05 |
| Proteoform-level cutoff type | FDR |
| Proteoform-level cutoff value | 0.05 |
| Allowed N-terminal forms | NONE, NME, NME_ACETYLATION, M_ACETYLATION |
| Maximum mass shift of modifications | 500 Da |
| Minimum mass shift of modifications | -500 Da |
| Thread number | 15 |
| E-value computation | Generation function |

**Table S2.** Hyperparameter settings for the FNN model

| Parameters | Search space | Selected value for RPLC | Selected value for CZE |
|---|---|---|---|
| #Hidden layer | [1,2,3] | 3 | 2 |
| Dense feature | [64,128,256,512,1024] | 128 | 256 |
| Dropout rate | [0,0.1,0.2] | 0 | 0 |

**Table S3.** Hyperparameter settings for the DeepRT+ model

| Parameters | Search space | Selected value for RPLC | Selected value for CZE |
|---|---|---|---|
| Filter size | [64,128,256] | 128 | 128 |
| Kernel size | [4,6,8,10,12,14,16] | 8 | 16 |
| Batch normalization | [0,1] | 0 | 0 |
| Batch size | (8,20) | 16 | 19 |
| #Epochs | [20,30,40] | 40 | 40 |

**Table S4.** Hyperparameter settings for the Prosit model

| Parameters | Search space | Selected value for RPLC | Selected value for CZE |
|---|---|---|---|
| Embedding dimension | [24,28,32] | 24 | 24 |
| GRU unit | [64,128,256,512] | 512 | 128 |
| Dense feature | [64,128,256,512] | 128 | 256 |

**Table S5.** Hyperparameter settings for the DeepDIA model

| Parameters | Search space | Selected value for RPLC | Selected value for CZE |
|---|---|---|---|
| Filter size | [16,32,64] | 64 | 16 |
| Kernel size | [3,4,5] | 4 | 3 |
| LSTM feature | [64,128,256] | 64 | 64 |
| Dense feature | [128,256,512] | 128 | 64 |

**Table S6.** Benchmarking of 5 machine learning models for proteoform retention time prediction on the LC-OT test data set. The models are trained using the LC-OT training set.

| Model | $R^2$ | $\Delta t_{r95\%}$ | MSE |
|---|---|---|---|
| GPTime | 0.837 | 0.488 | 0.00612 |
| FNN | 0.857 | 0.463 | 0.00535 |
| DeepRT+ | 0.771 | 0.522 | 0.00857 |
| Prosit | **0.860** | **0.486** | **0.00525** |
| DeepDIA | 0.792 | 0.552 | 0.00777 |

**Table S7.** Performance of the FNN model with 4 feature sets for migration time prediction on the CZE-SW480 data set with 5-fold cross validation. A total of 7 features are divided into 3 groups: (1) the molecular mass and the charge state, (2) the numbers of D, E, and N residues, and (3) the numbers of L and I residues.

| Feature groups | $R^2$ | MSE |
|---|---|---|
| 1 | 0.954 | 0.00040 |
| 1 and 2 | **0.959** | **0.00034** |
| 1 and 3 | 0.955 | 0.00040 |
| 1, 2, and 3 | 0.959 | 0.00035 |

**Table S8.** Benchmarking of 5 machine learning models for proteoform migration time prediction on the CZE-SW480 test data set. The models are trained using the CZE-SW480 training data set.

| Model | $R^2$ | $\Delta t_{r95\%}$ | MSE |
|---|---|---|---|
| Semi-empirical | 0.878 | 0.172 | 0.00118 |
| FNN | **0.945** | **0.120** | **0.00052** |
| DeepRT+ | 0.750 | 0.291 | 0.00243 |
| Prosit | **0.946** | **0.112** | **0.00052** |
| DeepDIA | 0.611 | 0.364 | 0.00282 |

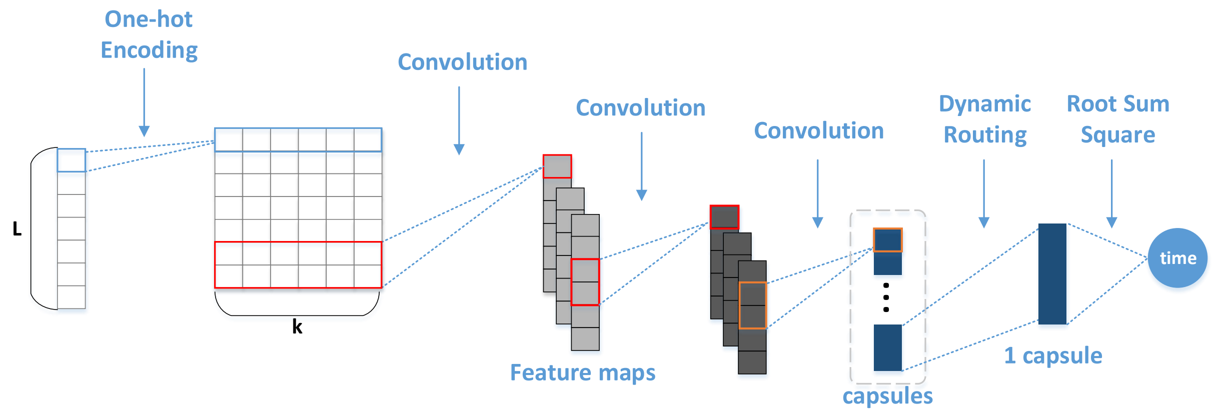


**Figure S1.** Architecture of the DeepRT+ model. The input protein sequence (length= $L$) is encoded with one-hot encoding to an $L \times k$ matrix where $L$ is the padded length (200) of the sequence and $k$ = 20 is the types of amino acid residues. The encoded matrix is fed into two convolutional layers followed by two capsule layers with dynamic routing. The root sum square (RSS) of the output vector of the final capsule is reported.

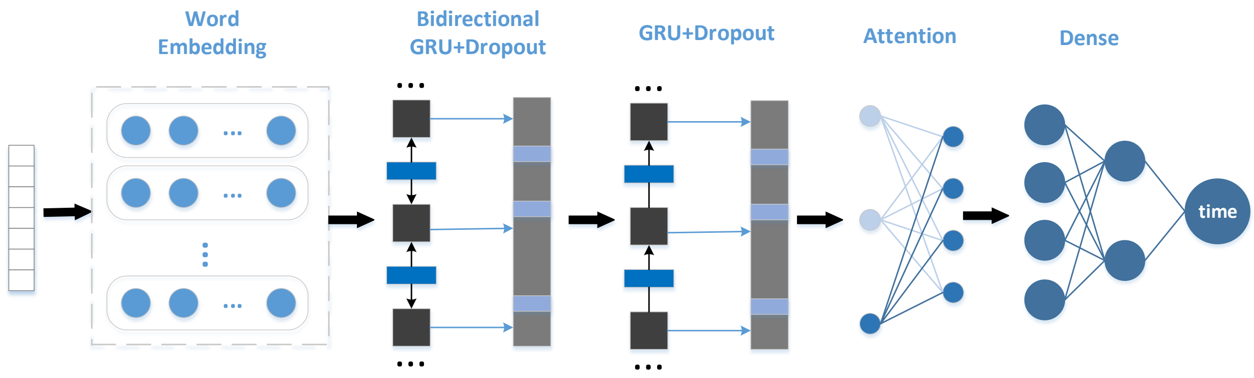


**Figure S2.** Architecture of the Prosit model. The input protein sequence is encoded by a word embedding layer, which is connected to one bidirectional recurrent layer with Gated Recurrent Unit (GRU) and one normal recurrent layer with GRU. The output from the dropout layer is flattened with an attention layer, which is connected to two dense layers.

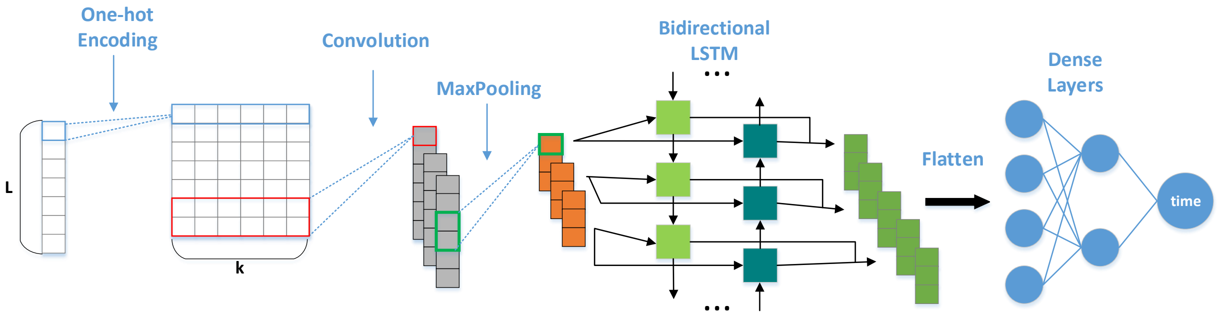


**Figure S3.** Architecture of the DeepDIA model. The input sequence (length= *L*) is encoded with one-hot encoding to an *L*×*k* matrix. The input features are fed into a convolutional layer with max pooling. A bidirectional LSTM layer is used to capture the sequential patterns in the output of max pooling.  The output of the bidirectional LSTM layer is flattened, and two dense layers are used to generate the final prediction result.
